# Transfer RNA levels are tuned to support differentiation during Drosophila neurogenesis

**DOI:** 10.1101/2024.09.06.611608

**Authors:** Rhondene Wint, Michael D. Cleary

## Abstract

Neural differentiation requires a multifaceted program to alter gene expression along the proliferation to differentiation axis. While critical changes occur at the level of transcription, post-transcriptional mechanisms allow fine-tuning of protein output. We investigated the role of tRNAs in regulating gene expression during neural differentiation by quantifying tRNA abundance in neural progenitor-biased and neuron-biased *Drosophila* larval brains. We found that tRNA profiles are largely consistent between progenitor-biased and neuron-biased brains but significant variation occurs for 10 cytoplasmic isodecoders (individual tRNA genes) and this establishes differential tRNA levels for 8 anticodon groups. We used these tRNA data to investigate relationships between tRNA abundance, codon optimality- mediated mRNA decay, and translation efficiency in progenitors and neurons. Our data reveal that tRNA levels strongly correlate with codon optimality-mediated mRNA decay within each cell type but generally do not explain differences in stabilizing versus destabilizing codons between cell types. Regarding translation efficiency, we found that tRNA expression in neural progenitors preferentially supports translation of mRNAs whose products are in high demand in progenitors, such as those associated with protein synthesis. In neurons, tRNA expression shifts to disfavor translation of proliferation-related transcripts and preferentially support translation of transcripts tied to neuron-specific functions like axon pathfinding and synapse formation. Overall, our analyses reveal that changes in tRNA levels along the neural differentiation axis support optimal gene expression in progenitors and neurons.

**AUTHOR SUMMARY:** Wint et al. quantified tRNA expression in *Drosophila* larval brains, using wildtype brains composed primarily of neurons and mutant brains composed primarily of neural progenitors (neuroblasts). These neuron-biased and neuroblast-biased tRNA measurements, combined with mRNA decay data and computational modeling of translation efficiency, revealed that: 1) tRNA abundance is largely constant between neural progenitors and neurons but significant variation exists for 10 nuclear tRNA genes and 8 corresponding anticodon groups, 2) tRNA abundance correlates with codon-mediated mRNA decay in neuroblasts and neurons but does not completely explain the different stabilizing or destabilizing effects of certain codons, and 3) changes in tRNA levels during differentiation shift translation optimization from a program supporting proliferation to a program supporting differentiation. These findings reveal coordination between tRNA expression and codon usage in transcripts that regulate neural development.

## INTRODUCTION

Neural differentiation requires regulation of gene expression at the transcriptional and post-transcriptional levels. One potential post-transcriptional mechanism involves enhancing or suppressing mRNA translation through changes in tRNA abundance. The degeneracy of the genetic code allows selection among synonymous codons in ways that enhance or repress translation, as determined by the abundance of cognate tRNAs. In tandem, tRNA expression may evolve to support optimal translation programs for a particular cell fate or physiologic condition. Far from serving as invariant components of the translation machinery, tRNAs are regulated in ways that affect protein output during developmental transitions, disease progression, and stress responses [1,2,3,4]. For example, in *S. cereviseae*, environmental stress induces expression of rare tRNAs that are necessary to translate codons enriched in stress-response mRNAs [5]. Similarly, mRNAs encoding proto-oncogenes are enriched in codons decoded by tRNAs that are more abundant in cancer and progenitor cells compared to differentiated cells [6].

Eukaryotic genomes typically contain multiple genes encoding tRNAs with the same anticodon and these tRNAs are known as isoacceptors. Individual tRNAs within an isoacceptor group are known as isodecoders and may be distinguished by sequence features outside the anticodon. The extent to which differential expression of tRNA genes (isodecoders) changes the anticodon pool for an amino acid (isoacceptor levels) is not straightforward and may be context dependent. An early analysis of tRNA and mRNA abundance across human tissues found tissue-specific differences in isoacceptor levels and these differences correlated with codon usage in highly expressed genes [7]. Subsequent work, using an alternative method for quantifying tRNAs, found that while there is significant variation in isodecoder expression across human tissues, isoacceptor levels remain largely unchanged [8]. This phenomenon has been termed “anticodon buffering”. Anticodon buffering was observed during the differentiation of human induced pluripotent stem cells into neural stem cells and neurons: transcription of individual tRNA genes changed during differentiation but isoacceptor levels remained constant [9]. In this case, anticodon buffering occurred via constitutive high-level expression of a core set of isodecoders within the anticodon groups. Similarly, cell type-specific tRNA profiling in the mouse brain identified differences in tRNA gene transcription that alter the isodecoder composition, but not the abundance, of anticodon groups [10].

Isoacceptor abundance is a major determinant of translation elongation rates. If a particular codon is used with high frequency across the transcriptome but the cognate charged tRNA is rare, ribosome translocation may slow or pause along certain transcripts. Such inefficiency across the transcriptome would negatively impact proteostasis and cell physiology. At the level of individual mRNAs, ribosome translocation is under selection to achieve rates that optimize co-translational protein folding and precise protein stoichiometry [11,12]. These parameters may be met via fine-tuned coordination of codon usage and tRNA abundance. In addition to translation efficiency, tRNA availability affects mRNA stability; codons decoded by abundant tRNAs (optimal codons) have a stabilizing effect on transcripts while codons decoded by rare tRNAs (non-optimal codons) have a destabilizing effect on transcripts [13]. This codon optimality-mediated decay (COMD) has been described in many organisms and model systems [14,15,16,17,18]. Relationships between codon usage, tRNA abundance, and mRNA stability in the nervous system are multifaceted. For example, tRNA modifications [19] and RNA-binding proteins [20, 21] influence COMD in neural cell types. Our prior work in *Drosophila* found that optimal codons have an attenuated effect on mRNA stability in the embryonic nervous system compared to whole embryos but this attenuation was not explained by the relative abundance of tRNAs that decode optimal codons [17].

Over 50 neurologic disorders have been linked to tRNA dysregulation [22], highlighting the importance of tRNAs in neural development and neural function. Here we used *Drosophila* larval neurogenesis as a model system to test the hypothesis that tRNA abundance and codon usage establish specific programs of mRNA stability and translation during neural differentiation. The *Drosophila* larval brain contains a population of neural progenitors, called neuroblasts, that undergo multiple rounds of asymmetric self- renewing divisions to produce a neuroblast and a differentiating cell. A genetic manipulation that locks neuroblasts into symmetric self-renewing divisions generates over 4,000 ectopic neuroblasts and few neurons per brain lobe [22]. In contrast, wildtype brains contain only 100 neuroblasts, roughly 250 glia, and approximately 5,000 neurons per brain lobe at the end of larval neurogenesis [23,24]. We and others have confirmed that the RNA from these two sources (ectopic neuroblast brains and wildtype brains) represent distinct “neuroblast-biased” and “neuron-biased” transcriptomes [23,25]. In the present study, we used this model system to quantify tRNA abundance in neuroblast-biased and neuron-biased brains and combined these data with measurements of mRNA abundance and mRNA decay in neuroblasts and neurons. This approach allowed us to interrogate relationships between tRNA levels, COMD and translation optimization during neural differentiation.

## RESULTS

### Neural progenitors and neurons have similar patterns of tRNA expression but variation exists for isodecoders and anticodon groups

To obtain tRNA profiles representative of proliferating neuroblasts, we used *insc-Gal4* to drive expression of *UAS-aPKC^CAAX^* in neuroblasts and harvested larval brains at 96 - 102 hours after larval hatching (ALH) as a source of neuroblast-biased RNA, as previously described [23,25]. We refer to data obtained from these samples as “neuroblast” data. In contrast, we used wildtype larval brains at 96 - 102 hours ALH as a source of neuron-biased RNA. We refer to data obtained from these samples as “neuron” data. These RNA samples were used for quantitative tRNA sequencing based on the hydro-tRNAseq method [26].

We obtained triplicate global tRNA profiles from neuroblast-biased and neuron-biased brains. First, we evaluated the abundance of tRNA isodecoders; tRNA genes that share the same anticodon but differ in their sequence body (not all isodecoders are distinguishable due to sequence redundancy). There are 295 cytosolic tRNA genes in the *Drosophila* nuclear genome: 289 encode tRNAs for the standard 20 amino acids, one encodes a selenocysteine tRNA, and five are pseudogenes [27]. 84 isodecoder groups are distinguishable by mature tRNA sequence (isodecoder groups may contain multiple copies of indistinguishable genes) and these tRNAs comprise 44 isoacceptor groups. We obtained measurements for all 84 cytosolic isodecoders and found that while expression was largely similar between neuroblasts and neurons, 10 tRNAs were differentially expressed (Figures 1A, 1B and supplemental table 1). These data also contained mitochondrial tRNA reads and two mitochondrial tRNAs were more abundant in neuroblasts (Figure 1B). For this study we chose to focus solely on cytosolic tRNAs (mitochondrial tRNA data are available in supplemental table 1). We measured the abundance of each cytosolic tRNA isoacceptor group by averaging the counts for all isodecoders in a group. 8 isoacceptor groups differed significantly: Pro-UGG, Pro-CGG, Gly-GCC, Glu-CUC, Tyr-GUA, Leu-CAA, and Arg-ACG were more abundant in neurons while Ser-UGA was more abundant in neuroblasts (Figure 1C).

**Figure 1.**
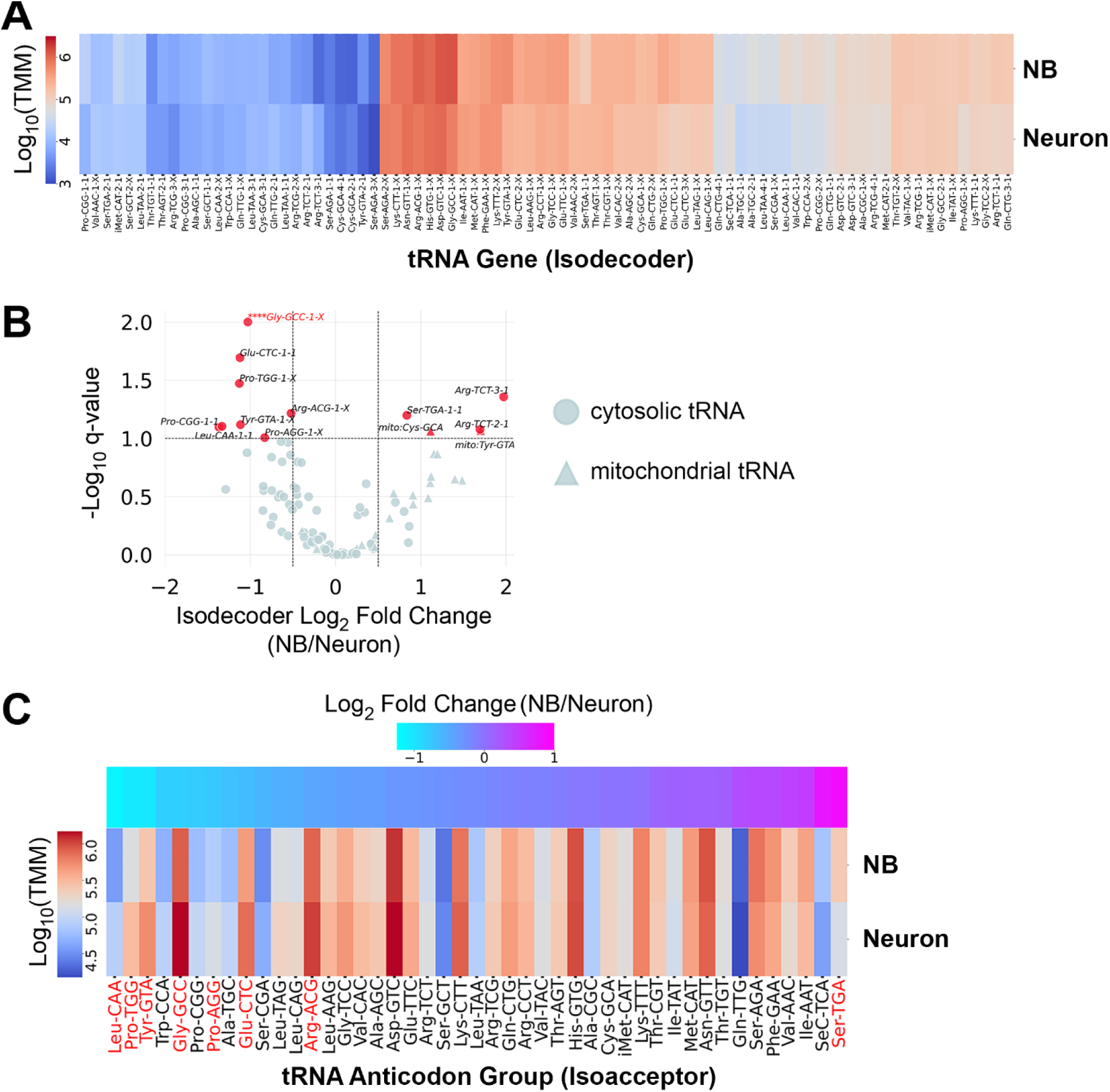
tRNA levels are largely constant between neuroblast-biased and neuron-biased brains, but variation exists for a subset of anticodon groups. (A) Hierarchically clustered heatmap showing the average expression level of 84 nuclear-encoded Drosophila tRNA genes (isodecoders). Counts were normalized to Trimmed Mean of M-values (TMM). An X in the isodecoder name indicates that more than one gene encodes that particular isodecoder and those genes are not distinguishable by mature tRNA sequence. (B) Volcano plot showing the relative isodecoder expression level for nuclear-encoded (cytosolic) and mitochondrial tRNA genes on the x-axis (fold change = average Neuroblast TMM / average Neuron TMM) and the false discovery rate q-value. tRNAs with a log2 fold change ≥ ± 0.5 and q-value ≤ 0.1 are highlighted in red. **** denotes that Gly-GCC-1-X is off scale, with a q-value < 0.001. (C) Hierarchically clustered heatmap showing the average expression level (normalized to Trimmed Mean of Mvalues (TMM)) for 44 anticodon groups (isoacceptors). Differentially expressed isoacceptors (q-value < 0.1) are highlighted in red.

### tRNA abundance correlates with codon optimality-mediated mRNA decay within neuroblasts and neurons

We used recently reported neuroblast-specific and neuron-specific mRNA decay measurements [25] to calculate codon stabilization coefficients (CSCs) for each cell type. As previously described, CSC is the Pearson’s coefficient between codon frequency in a transcript and transcript stability [14]. Codons identified as stabilizing (positive CSC) and codons identified as destabilizing (negative CSC) were largely the same in neuroblasts and neurons (Figure 2A). To investigate how COMD might regulate functionally related transcripts in the developing brain, we first identified 5,169 transcripts that are expressed in both neuroblast-biased and neuron-biased brains. Next we ranked these transcripts based on neuroblast- defined optimal or non-optimal codon content and performed gene ontology (GO) analysis on the top optimal codon-enriched transcripts and the top non-optimal codon enriched transcripts (Figure 2B).

**Figure 2.**
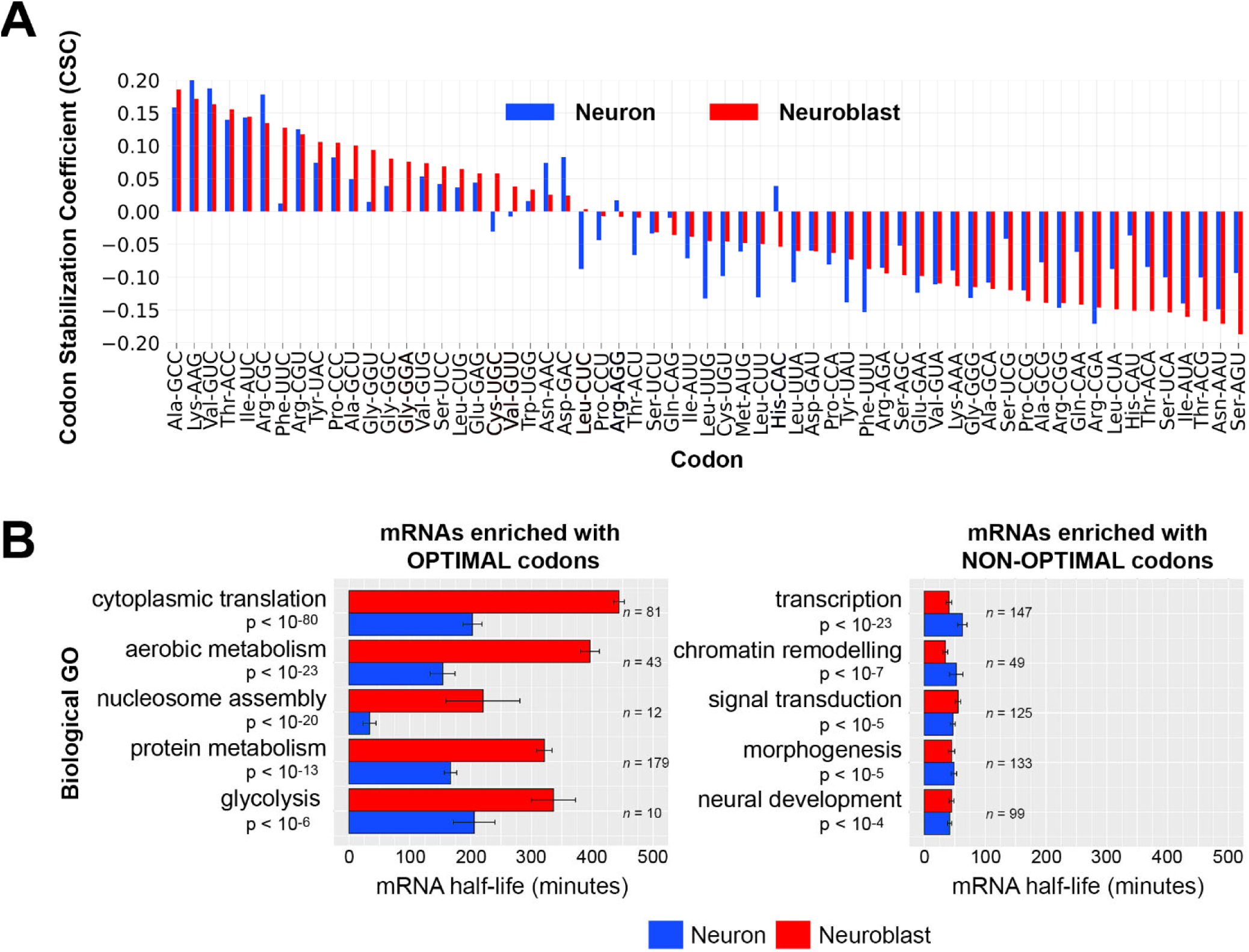
**Codon optimality-mediated decay is similar in neuroblasts and neurons and correlates with the stability of functionally related mRNAs.** (A) Codon stabilization coefficients (CSCs) were calculated using neuroblast-specific and neuron-specific transcriptome-wide mRNA decay measurements [24]. (B) Gene ontology (GO) categories of mRNAs enriched with optimal codons or non-optimal codons. 5,169 neural transcripts (mRNAs present in neuroblast-biased and neuron-biased brains) were ranked by percent optimal codon content (defined by optimality in neuroblasts): the top 10% optimal codon enriched mRNAs (left panel) and the top 10% non-optimal codon enriched mRNAs (right panel) were used for GO analysis. The top five (ranked by adjusted p-value) non-redundant GO categories and corresponding adjusted p-values are listed. Neuroblast-specific and neuron-specific mRNA decay data [24] for the transcripts within each GO category were used to calculate average mRNA half-life. The number of transcripts in each GO category is listed to the right of the plot bars. Error marks are standard error of the mean. Adjusted p-value is based on the Bonferroni correction.

Optimal codon enriched transcripts were significantly associated with metabolic processes that are elevated in proliferating cells, such as “cytoplasmic translation” and “aerobic metabolism”. Transcripts in these categories were more stable in neuroblasts than in neurons, even though 19 of the 22 enriched codons are optimal (CSC > 0.01) in both cell types. Non-optimal codon enriched transcripts were significantly associated with conditional cellular processes, such as “transcription”, “signal transduction” and “neural development”. Transcripts in these categories had similarly short half-lives in neuroblasts and neurons.

To investigate how tRNA abundance might explain CSCs within each cell type, we first compared tRNA isoacceptor abundance and CSCs based on Watson-Crick base pairing. These analyses revealed that, in both cell types, stabilizing codons tend to have abundant cognate tRNAs while destabilizing codons tend to have less abundant cognate tRNAs (Figure 3A). These results agree with the general model of how tRNA abundance determines the stabilizing or destabilizing effects of codons. A limitation of only using Watson-Crick base pairing is that several codons are wobble translated by multiple tRNAs. To overcome this limitation, we computed each codon’s tRNA adaptation index (tAI). tAI is an indicator of a codon’s translation efficiency that incorporates tRNA abundance and wobble interactions [28]. tAI calculations traditionally use tRNA gene copy number as a proxy for tRNA abundance, however we were able to use our tRNA measurements to calculate neuroblast-biased tAI (NB-tAI) and neuron-biased tAI (neuron-tAI). In addition, we normalized tAIs within each amino acid family so that the most optimal synonymous codons (codons with the greatest decoding potential based on tRNA abundance) have a tAI equal to 1.0. Using these tAI values, we found a significant association between a codon’s tRNA- dependent translation efficiency (tAI) and its stabilizing influence on mRNA half-life (Figure 3B).

**Figure 3.**
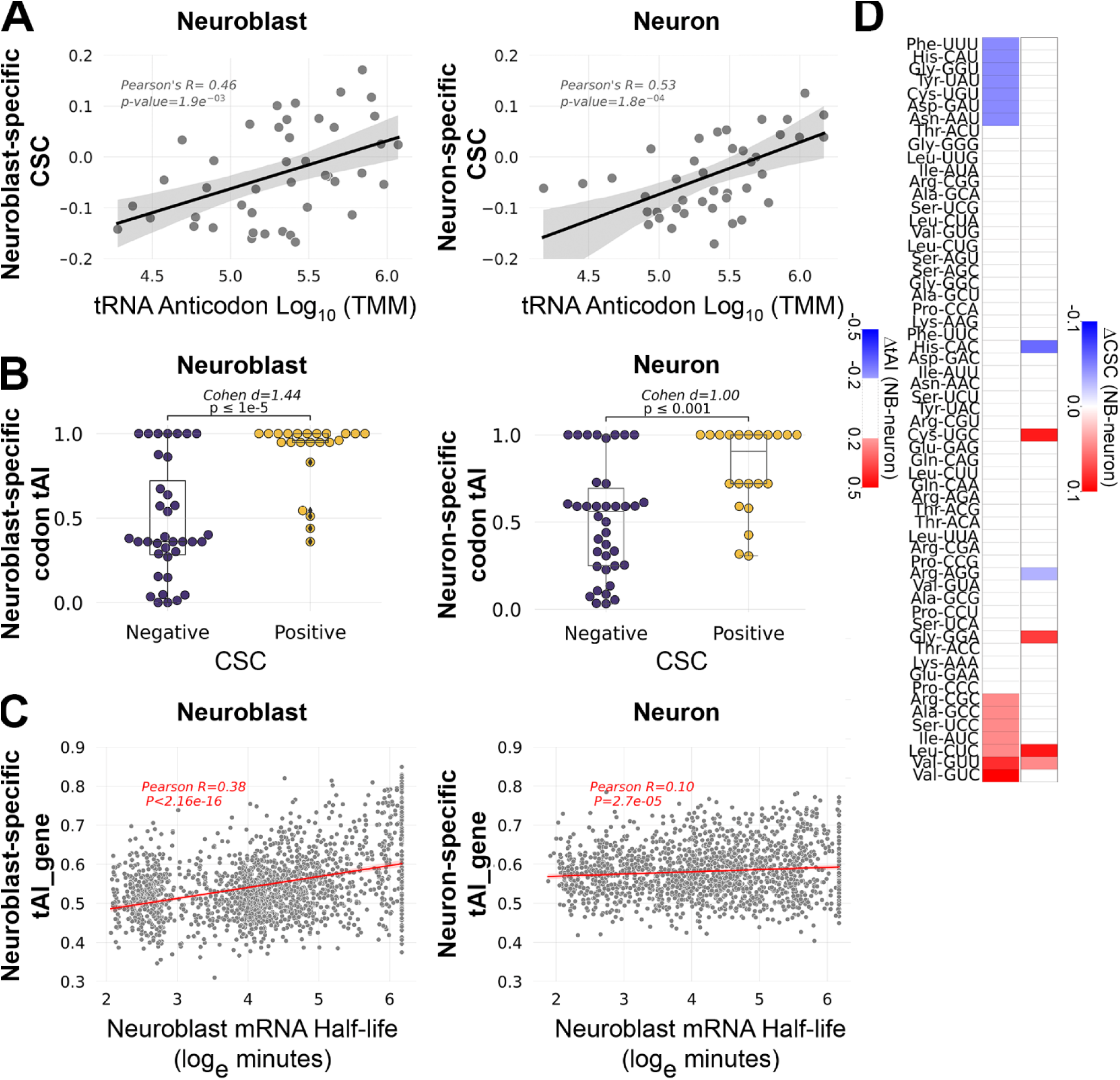
**tRNA abundance correlates with codon-mediated mRNA decay in neuroblasts and neurons.** (A) Scatter plots comparing tRNA isoacceptor abundance and codon stabilization coefficient (CSC) in each cell type (neuroblast data in left panel, neuron data in right panel). (B) Relationship between tRNA adaptation index (tAI) and CSC in each cell type (neuroblast data in left panel, neuron data in right panel). The effect size was measured by Cohen’s d and significance (p-value) was determined by Welch’s t-test. (C) Scatterplot comparing tAI_gene and mRNA half-life for 5,169 transcripts present in neuroblast-biased and neuron-biased brains. (D) Heatmaps comparing differential tAI (left column) and differential CSC (right column). Only absolute Δ tAI values ≥ 0.2 are shown. Only CSCs that differed in category (optimal, neutral, non-optimal) are shown, according to the following criteria: optimal (CSC ≥ 0.01) in one cell type but neutral (0.01 > CSC > -0.01) or non-optimal (CSC ≤ -0.01) in the other.

However, there is a stronger association between tAI and CSC in neuroblasts (Cohen’s d=1.44) than in neurons (Cohen’s d=1.00), with nearly twice as many optimal codons with a tAI of 1.0 in neuroblasts (18 optimal codons) than there are in neurons (11 optimal codons). This difference suggests that for common neural transcripts (mRNAs present in neuroblasts and neurons), tRNA adaptation is likely to have a stronger stabilizing effect in neuroblasts than in neurons. To test this prediction, we used the 5,169 transcripts that are expressed in both neuroblast-biased and neuron-biased brains and calculated the geometric mean of the tAI values for all codons in each transcript, a value we call tAI of a gene (tAI_gene). This revealed a stronger correlation between mRNA half-life and tAI_gene in neuroblasts than in neurons (Figure 3C). Overall, these analyses revealed a relationship between tAI and half-life, as expected, but neuron-tAI is a weaker predictor of transcript stability than NB-tAI.

Given that tAI is generally predictive of the stabilizing or destabilizing effects of codons within neuroblasts or neurons, we next asked if differential tAI is predictive of differential CSC. Only 6 codons had divergent CSC values in which the codon was optimal (CSC ≥ 0.01) in one cell type but neutral (0.01 > CSC > - 0.01) or non-optimal (CSC ≤ -0.01) in the other (Figure 3D). For the majority of codons analyzed (41 codons), tAI and CSC do not significantly differ between neuroblasts and neurons (Figure 3D). In most other cases there is no clear relationship between tAI and CSC: four codons have differential CSC but no difference in tAI and 12 codons have differential tAI but no difference in CSC. However, Leu-CUC and Val-GUU have a direct relationship between tAI and CSC, with increased tAI in neuroblasts corresponding to increased CSC in neuroblasts (Figure 3F). These results suggest that while tRNA abundance influences COMD within neuroblasts or neurons, tRNA abundance is a less robust predictor of differential COMD, with the exception of tRNAs that decode Leu-CUC and Val-GUU. We preformed GO analysis to see if transcripts enriched in CUC and/or GUU codons encode proteins with a common function. Compared to the set of 5,169 neural transcripts, mRNAs enriched (≥ 2% of codons) in CUC alone or GUU alone did not comprise any significant GO category. However, transcripts enriched in either of these codons (≥ 2% CUC alone, CUC and GUU, or GUU alone) yielded a single significantly enriched GO category: “Nucleosome Assembly” (Bonferroni adjusted p-value = 3.3 x 10^-7^). This result is due to abundant representation of replication-dependent histone mRNAs (H2A, H2B, H1, H3 and H4) among the CUC- and/or GUU-enriched transcripts. Replication-dependent histone mRNAs are significantly more stable in neuroblasts compared to neurons, as shown for the “Nucleosome Assembly” GO category in Figure 2B. These results suggest that decreased abundance of Leu-CUC and Val-GUU tRNAs contributes to the decreased stability of replication-dependent histone mRNAs in neurons.

### tRNA adaptation supports neuroblast-specific versus neuron-specific translation programs

We next asked if changes in tRNA abundance during neural differentiation support a shift from translation of mRNAs involved in progenitor-specific functions to translation of mRNAs involved in neuron-specific functions. Hierarchical clustering of the 5,169 common neural transcripts according to neuron-tAI_gene and NB-tAI_gene revealed several patterns of translation adaptation during neural differentiation (Figure 4A). We performed gene ontology analysis on the following tAI_gene clusters: a “shared high” group with high translation adaptation in both cell types, a “shared low” group with low translation adaptation in both cell types, and a “neuron up” group with increased translation adaptation in neurons relative to neuroblasts (Figure 4B). This revealed distinct biological functions associated with transcripts in each group. The “shared high” group is enriched in transcripts that support housekeeping processes important to neuroblasts and neurons, such as nuclear transport and protein transport. The “shared low” group is uniquely enriched in transcripts supporting conditional processes that occur in neuroblasts and newborn neurons, such as cell fate specification, Notch signaling and EGFR signaling. The “neuron up” group is uniquely enriched in transcripts supporting the neuron-specific process of synaptic transmission as well as more general processes like RNA splicing and protein kinase activity.

**Figure 4.**
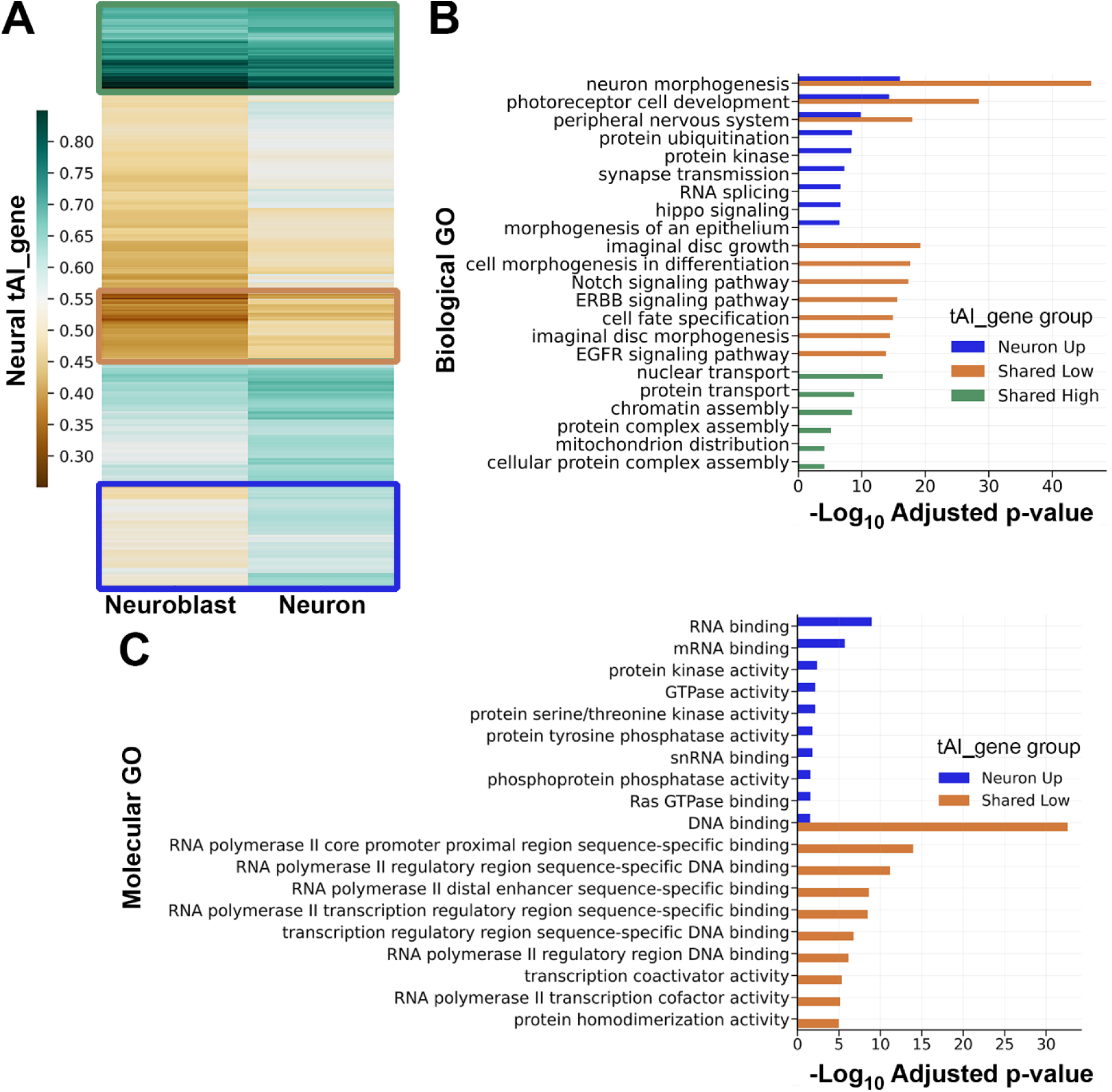
**tRNA adaptation during neural differentiation supports distinct translation programs.** (A) Hierarchically clustered heatmap showing neuroblast-specific and neuron-specific tAI_gene values for 5,169 neural mRNAs. Three emergent patterns are outlined in colored boxes: green for high tAI_gene in neuroblasts and neurons (“shared high”, n = 517 mRNAs), tan for low tAI_gene in neuroblasts and neurons (“shared low”, n = 411 mRNAs), and blue for increased tAI_gene in neurons compared to neuroblasts (“neuron up”, n = 965 mRNAs). (B) Biological function gene ontology (GO) category enrichment in each of the tAI_gene groups identified in part A. Bar color matches the group color in part A. Up to ten of the most significantly enriched GO categories, with adjusted p-value ≤ 0.0001, are listed (fewer categories if the p-value cutoff was not met). Adjusted pvalues are based on the Bonferroni correction. (C) Molecular function GO category enrichment for transcripts present in the “neuron morphogenesis”, “photoreceptor cell development”, and “peripheral nervous development” Biological GO categories shown in part B. The ten most significantly enriched Molecular GO categories for each group (neuron up, shared low) are listed. Adjusted p-value is based on the Bonferroni correction.

The “shared low” and “neuron up” groups both yield the GO categories “neuron morphogenesis”, “photoreceptor cell development”, and “peripheral nervous development”. In these overlapping GO categories, “shared low” transcripts (poorly adapted to translation in neurons) overwhelmingly encode transcription factors while “neuron up” transcripts (well adapted to translation in neurons) overwhelmingly encode kinases, small GTPases and RNA binding proteins (Figure 4C).

### mRNAs enriched in proliferation-associated codons are less efficiently translated by the post- differentiation tRNA pool

To further investigate the relationship between codon usage and tRNA abundance during neural differentiation, we used relative codon frequencies across the 5,169 neural transcripts to represent each coding sequence as a 59-dimensional vector. This allowed us to visualize neural transcriptome codon distribution via principal component analysis (PCA) followed by k-means unsupervised clustering. PCA on codon usage captured a total variance of 23%, with PC1 covering 15% of the variation and PC2 covering 8% of the variation (**Figure 5A**). Because PC1 captured more of the variation, we focused on mRNAs belonging to clusters that project to opposite poles of PC1: cluster 0 and cluster 4. GO analysis revealed that cluster 0 mRNAs (*n*=373) are significantly associated with proliferation-related terms such as cytoplasmic translation and mitochondrial metabolism, while cluster 4 mRNAs (*n*=219) are more significantly associated with GO terms related to neural differentiation and neuron function (**Figure 5B**). Based on this distribution of mRNAs supporting distinct biological functions, we assigned cluster 0 a “Proliferation” identity and cluster 4 a “Differentiation” identity.

**Figure 5.**
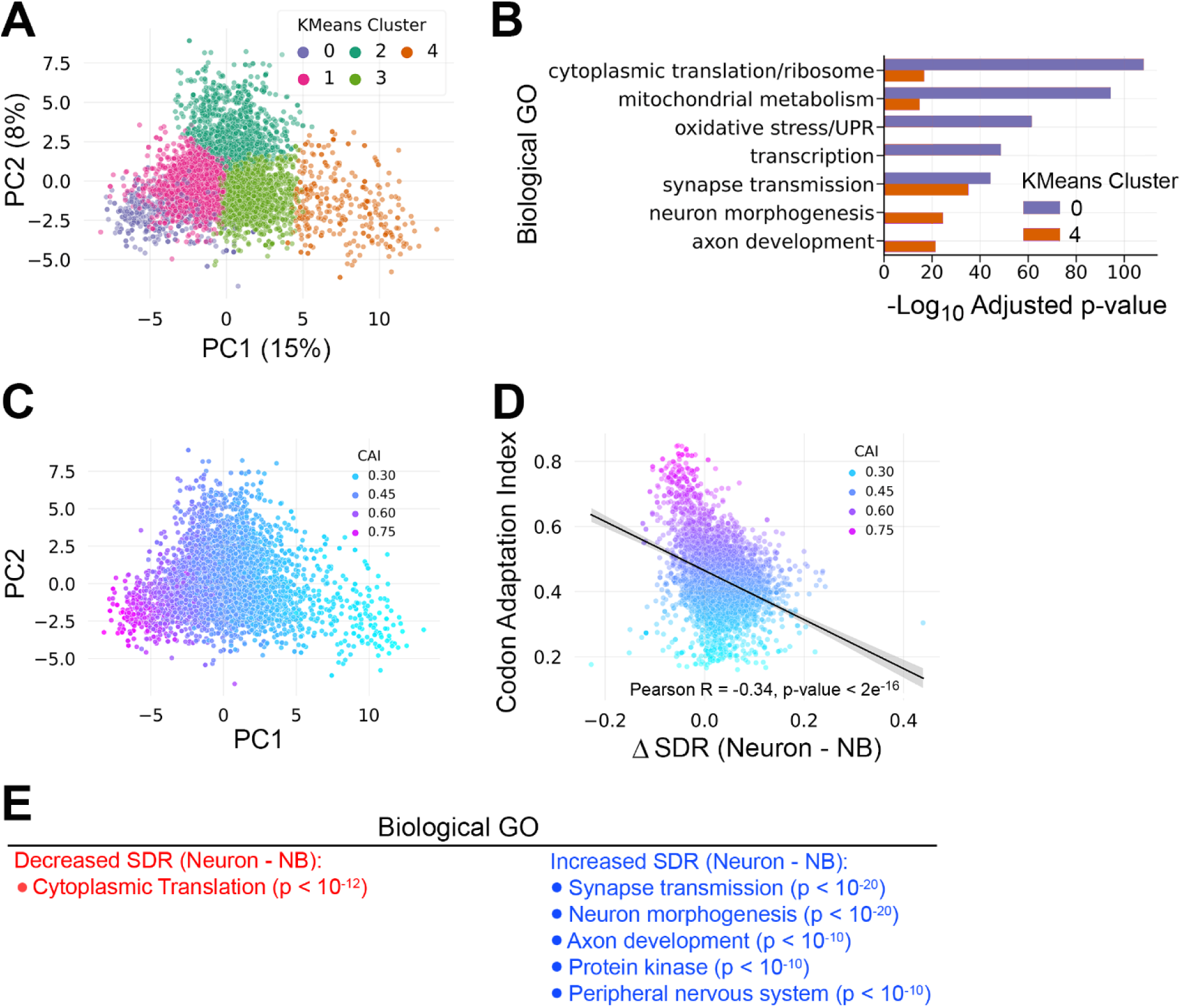
**mRNAs enriched in proliferation-associated codons are poorly adapted for translation by the post-differentiation tRNA pool.** (A) Principal component analysis (PCA) on the normalized codon frequencies of 5,169 mRNAs present in neuroblast-biased brains and neuron-biased brains. PCA captured a total variation of 23%, with 15% of the variance in principal component 1 (PC1). Unsupervised clustering by kmeans grouped the mRNAs into 5 clusters. (B) GO analysis of mRNAs within kmeans clusters at opposite ends of PC1: cluster 0 and cluster 4. (C) PCA plot from part A recolored with the Codon Adaptation Index (CAI) of each mRNA. (D) Scatterplot comparing CAI and the change in Supply Demand Ratio (SDR) between neurons and neuroblasts (ΔSDR = Neuron SDR – Neuroblast SDR). The Pearson R value and line of best fit show the inverse relationship between CAI and neuron-optimal SDR. (E) The 5,169 neural mRNAs were ranked by _SDR (Neuron SDR – Neuroblast SDR) and the top 10% (increased SDR) and bottom 10% (decreased SDR) were subject to GO analysis. Significantly enriched GO categories with a Bonferroni adjusted p-value < 10-5 are listed.

Next, we calculated the codon adaptation index (CAI) for transcripts in all five k-means clusters. CAI quantifies the similarity between a gene’s synonymous codon usage and codon usage in a reference set of highly expressed genes, typically ribosomal protein genes [29]. We used *Drosophila* ribosomal protein genes as the reference for CAI calculations and recolored the PCA plot according to CAI. This analysis revealed that CAI decreases along the PC1 axis, from the Proliferation Cluster (cluster 0) to the Differentiation Cluster (cluster 4) (Figure 5C). This inverse relationship between CAI and differentiation suggests that transcripts associated with differentiation are biased against inclusion of codons that are optimally translated during proliferation.

To further model how the intersection of tRNA abundance and mRNA expression leads to changes in translation efficiency during neural differentiation, we calculated the supply – demand ratio for each codon-anticodon pair (SDR_c_) [11]. Previous work has shown that SDR is a better predictor of translation efficiency than tAI alone [30]. Here SDR_c_ is the ratio between tAI (the supply of tRNAs) and the frequency of the corresponding codon weighted by mRNA abundance (the demand for cognate tRNAs) in either neurons or neuroblasts. We used SDR_c_ values to calculate transcript-level SDR by taking the geometric mean of SDR_c_ values across each coding sequence. SDR therefore integrates static codon frequency information with dynamic tRNA and mRNA abundance information.

This approach allowed us to measure how the SDR of individual transcripts differs between neurons and neuroblasts (ΔSDR = neuron SDR – neuroblast SDR). In this case, a positive ΔSDR indicates improved translation post-differentiation. ΔSDR values ranged from -0.23 (worst adapted to neuron translation) to 0.44 (best adapted to neuron translation), with a median of 0.033 (Figure 5D). Since we previously identified proliferation-associated transcripts as having a high CAI, we asked how this sequence-intrinsic property (codon adaptation) correlates with translation efficiency as defined by SDR. We found a significant negative correlation between ΔSDR and CAI (Figure 5D), indicating that mRNAs enriched in proliferation-associated codons are less efficiently translated by the post-differentiation tRNA pool. To assess the biological relevance of SDR changes during differentiation, we performed GO analysis on mRNAs within the top and bottom 10% of ΔSDR values. mRNAs with decreased SDR upon differentiation are uniquely associated with cytoplasmic translation (Figure 5E). In contrast, mRNAs with increased SDR upon differentiation are uniquely associated with post-mitotic functions like synaptic transmission and neuron morphogenesis (Figure 5E).

## DISCUSSION

The extent to which differential tRNA availability regulates cell type-specific gene expression is an important question in developmental biology. An mRNA may be present in a progenitor cell and its differentiated progeny, but the stability and translation of that mRNA depends on the relative abundance of tRNAs that decode the message. Here we investigated the role of tRNAs in shaping gene expression within the context of *Drosophila* neural development. We identified individual tRNA genes that are differentially expressed between neuroblasts and neurons and in all but one case this differential isodecoder expression is sufficient to establish neuroblast-specific and neuron-specific anticodon pools. As expected based on the known mechanism of codon optimality-mediated mRNA decay (COMD), tRNA anticodon abundance strongly influences mRNA stability within each cell population. However, tRNA abundance only partially explains differences in the stabilizing or destabilizing effects of certain codons between neuroblasts and neurons, suggesting that additional factors affect cell type-specific mRNA decay. We identified a more straightforward relationship between tRNA anticodon abundance and translation efficiency. Across the neural transcriptome, the tRNA profile of neuroblasts is highly adapted to support translation of proliferation and growth-associated transcripts while the tRNA profile of neurons is highly adapted to support translation of mRNAs associated with axonogenesis and synapse function. These findings provide insights into the role of differential tRNA expression in dynamic regulation of protein synthesis during neural differentiation.

Hydro-tRNAseq proved to be an effective method of quantifying tRNA abundance in neuroblast-biased and neuron-biased brains. We identified all 84 isodecoders that are distinguishable by mature tRNA sequence at counts spanning several orders of magnitude (1x10^6^ to over 1x10^11^ transcripts per million mapped reads). These data revealed significant variations in the expression of ten isodecoders. In the majority of cases (8/10), differential expression of a single isodecoder was sufficient to establish a corresponding difference in the anticodon pool. This is because the differentially expressed isodecoders are present at much higher levels than the constitutively expressed tRNAs of the same anticodon group. For example, in the tRNA:Ser-UGA group (composed of two genes), tRNA:Ser-UGA-1-1, which is more abundant in neuroblasts, is expressed at high levels in both cell populations (220,526 trimmed mean of M-values (TMM) in neuroblasts, 123,522 TMM in neurons) while the constitutively expressed tRNA:Ser- UGA-2-1 is expressed at low levels in both cell populations (14,398 TMM in neuroblasts, 12,522 TMM in neurons) (Figure 1A and supplemental table 1). The inverse is observed for the two differentially expressed tRNA:Arg-UCU isodecoders that did not support differential anticodon abundance. tRNA:Arg- UCU-2-1 (12,426 TMM in neuroblasts, 3,840 TMM in neurons) and tRNA:Arg-UCU-3-1 (6,983 TMM in neuroblasts, 1,779 TMM in neurons) are more abundant in neuroblast-biased brains but are expressed at low levels compared to the constitutively expressed tRNA:Arg-UCU-1-1 (92,728 TMM in neuroblasts, 141,178 TMM in neurons). This is the only example where we observed anticodon buffering; all other differential isodecoder expression caused corresponding differences in isoacceptor pools. This is in contrast to the widespread anticodon buffering observed across human tissues [8], in an *in vitro* model of neural differentiation [9], and among differentiated cells of the mouse brain [10].

Why is anticodon buffering less common in the developing *Drosophila* brain? One potential explanation is that there are fewer tRNA genes per anticodon group in *Drosophila* compared to humans. This makes it less likely that a cohort of isodecoders will buffer the differential expression of a single tRNA gene. For example, the *Drosophila* tRNA:Ser-UGA anticodon group is composed of only two genes but there are five tRNA:Ser-UGA genes in humans. With respect to neural differentiation, another potential explanation is that *Drosophila* neurogenesis occurs over the course of hours instead of the weeks required to differentiate human iPS cells to neurons *in vitro* [9] or the similarly prolonged timeframe of *in vivo* mammalian neurogenesis [31]. Rapid neurogenesis might select for post-transcriptional mechanisms to reinforce necessary patterns of gene expression, particularly to limit translation of aberrantly expressed transcripts or progenitor-specific transcripts passed to differentiating progeny via cytoplasmic inheritance. Such rapid and dynamic changes in tRNA anticodon pools can only occur in the absence of anticodon buffering.

Our quantitative tRNA data allowed several lines of inquiry. First we set out to investigate the relationship between anticodon abundance and COMD. We discovered the expected relationship between CSC and tRNA levels within each cell population, with correlations similar to those reported for other systems [18]. We evaluated standard Watson-Crick base pairing between codons and anticodons in addition to wobble interactions that allow a codon to be read by more than one tRNA, using the tRNA adaptation index (tAI). Our tAI analysis had the advantage of incorporating tRNA measurements as opposed to relying on tRNA gene copy number as a proxy for tRNA abundance. We were interested to discover that tAI and tAI_gene are less robust predictors of codon optimality and transcript stability in neurons compared to neuroblasts (Figure 3B and C). This is reminiscent of the attenuated COMD we previously described in the embryonic nervous system [17].

The weaker relationship between tRNA abundance and mRNA decay in neurons may explain why differential tAI only occasionally corresponds to differential CSC. It may be that mRNA decay in neurons is more sensitive to orthogonal decay mechanisms, with *trans*-acting factors complimenting or counteracting the effects of COMD. This may be similar to how the RNA-binding protein FMRP preferentially binds and stabilizes optimal codon-containing transcripts in mouse neurons [20] or how the miR-430 microRNA antagonizes the stabilizing effects of optimal codons during zebrafish embryogenesis [32]. One likely regulator of such *trans*-acting attenuation of COMD is Orb2, an RNA-binding protein that was recently found to preferentially stabilize non-optimal codon containing transcripts in *Drosophila* neurons [21]. An additional explanation for the lack of correlation between differential CSC and differential tRNA in some cases may be that the hydro-tRNAseq method does not distinguish between charged (amino acid attached) and uncharged (no amino acid) tRNAs. It is possible that the differential CSCs we identified are established by unequal charging of the corresponding tRNAs in neuroblasts versus neurons. Regardless, we did identify two codons with corresponding differences in tAI and CSC. Those codons (Leu-CUC and Val-GUU) are significantly over-represented in transcripts encoding replication-dependent histones that are only transcribed during S-phase. Decreased abundance of tRNAs that decode CUC and GUU in neurons may contributes to COMD of aberrantly expressed histone transcripts while the increased abundance of those tRNAs in neuroblasts may ensure efficient production of histones necessary to support DNA replication. Replication-dependent histone mRNAs lack poly-A tails and have a truncated 3’ UTR that forms a stem-loop structure [33]. These features could make replication-dependent histone mRNAs particularly sensitive to COMD since common 3’ UTR targeting mechanisms are not expected to influence codon-mediated effects on stability.

We used multiple metrics to investigate the impact of tRNA abundance on translation efficiency. Our first metric, gene-level analysis of tRNA adaptation (using tAI_gene), revealed that neuroblast-specific and neuron-specific tRNA pools establish distinct translation programs (Figure 4). For example, many transcripts supporting early stages of neurogenesis (such as those in the “neuron morphogenesis” GO category) have low translation adaptation in neuroblasts and neurons. This may reflect a need for low-level synthesis of these proteins in progenitors so that appropriate amounts are immediately present (via cytoplasmic inheritance) in newborn neurons. In contrast, transcripts that support functions unique to mature neurons (in particular those in the “synapse transmission” GO category), have a marked increase in neuron-tAI_gene relative to neuroblast-tAI_gene, as expected for proteins that serve no purpose in the initial stages of neurogenesis. Another dichotomy is revealed when transcripts with differential tAI_gene are classified by protein function (figure 4C). Transcription factors have low tAI_gene in neuroblasts and neurons while RNA-binding proteins and kinases have a neuron-specific increase in tAI_gene. This distinction suggests that transcription factors, particularly those that initiate cell fate decisions, are relatively inefficiently translated in order to ensure transient activity while downstream effectors of neuronal processes (RNA-binding proteins, kinases, small GTPases) have increased translation efficiency to ensure sustained protein abundance.

As a second translation efficiency metric, we used PCA to identify clusters of proliferation-associated and differentiation-associated transcripts and determined the codon adaptation index (CAI) of those transcripts (Figures 5A – C). CAI measures how well a transcript’s codon usage aligns with the codon usage of a reference set of highly expressed proteins. We followed standard convention and used transcripts encoding ribosomal proteins as the reference set. This revealed that proliferation-associated transcripts have a high CAI while CAI decreases in differentiation-associated transcripts. High CAI in proliferation-associated transcripts might be expected since this group contains many ribosomal protein mRNAs, but decreased CAI in differentiation-associated transcripts is informative. We and others have previously shown that neural progenitors have higher rates of protein synthesis [34,35] but there is no *a priori* reason that post-differentiation transcripts would not employ the same set of codons found in ribosomal proteins to ensure efficient translation. The fact that transcripts supporting neuron-specific functions avoid proliferation-associated codons suggests that a primary function of differential tRNA abundance during *Drosophila* neurogenesis is to shift optimal translation toward a unique set of differentiation-associated transcripts.

To test the above prediction, we employed a third translation efficiency metric: the supply – demand ratio (SDR). SDR provides a measure of effective tRNA levels by normalizing anticodon abundance to codon demand across the transcriptome. Here we had the advantage of pairing neuroblast-biased tRNA measurements with neuroblast-biased mRNA abundance and neuron-biased tRNA measurements with neuron-biased mRNA abundance. This SDR analysis further confirmed that the translation efficiency of proliferation-associated transcripts decreases upon differentiation while the translation efficiency of neuron-associated transcripts increases (Figure 5D). Decreased SDR upon neural differentiation is expected to significantly dampen production of ribosomal proteins and other regulators of cytoplasmic translation (Figure 5E). This result supports a model in which the drop in protein synthesis that occurs during neural differentiation is driven, at least in part, by decreased translation of transcripts encoding components of the translation machinery. Conversely, increased SDR upon neural differentiation is expected to enhance the translation of transcripts associated with neuron maturation and function, particularly synaptic transmission and neuron morphogenesis (Figure 5E). It is noteworthy that our SDR analysis (incorporating anticodon and codon abundance) recapitulates GO results obtained using codon bias alone (independent of any anticodon data) (compare Figures 5B and 5E). This suggests that tRNA levels can strongly influence translation programs during neural differentiation, independent of changes in mRNA abundance.

Transcription is not the sole determinant of gene expression during neurogenesis. For example, time- course analysis of human ESC differentiation into neurons revealed that mRNA levels failed to completely explain the observed variance in protein expression [36]. Our work suggests that changes in tRNA expression establish translation programs that meet the distinct needs of neuroblasts and neurons. We propose that the shift away from optimal translation of progenitor-associated transcripts and toward optimal translation of differentiation-associated transcripts is a form of tRNA driven “canalization” [37] which reinforces commitment to neural differentiation.

## MATERIALS AND METHODS

### Drosophila Genetics and larval brain preparation

The Bloomington Drosophila Stock Center supplied Oregon-R-P2 (a wildtype strain, stock # 2376) and *insc-Gal4* (stock # 8751) lines. The *UAS-aPKC^CAAX^* line was provided by C.Y. Lee. Flies were raised at 23-25°C. Whole brains were dissected from late-stage larvae (120 hours after larval hatching) of Oregon-R-P2 and transgenic ectopic-neuroblast mutant (*UAS-aPKC^CAAX^* x *Insc-Gal4*) flies. A total of 3 biological replicates were obtained for each genotype.

### Hydro-tRNAseq library preparation

We used methods described by Gogakos et al. [25] to generate hydro-tRNAseq libraries.

### Bioinformatics and statistics

#### Hydro-tRNAseq data analysis

Raw sequencing reads were preprocessed with fastqc, and quality-trimmed using cutadapt (v3.7) to remove adaptor sequences and reads shorter than 12bp. Custom reference *D.melanogaster* tRNA transcriptome was generated by collapsing identical mature tRNA sequences from gtRNAdb (UCSC genome release 6.40) and adding a CCA at the 3’ end of each sequence. tRNAseq reads were mapped to the reference tRNA transcriptome using subread-align (v.2.01) [38] with the following parameters, *-t 1 - T 64 --multiMapping -B 5 -m 3 -I*, to allow up to 3 mismatches and report the first 5 multi-mapped reads. For quality control, only aligned reads with MAPQ>=10 were retained. Read count summary was performed using subread’s featureCounts, in which multi-mapped reads were fractionally split between their references. NOISeqBio in R(v.3.6) was used to perform read count normalization (TMM), batch correction, and differential gene expression analysis. NOISeqBio uses non-parametric Bayesian methods to estimate differentially expressed genes in biological replicates and reports a q-value that is statistically equivalent to the Benjamini-Hochberg false-discovery rate.

#### Larval brain mRNA abundance and mRNA half-life measurement

RNA abundance and mRNA half-life values were calculated using data from Sami et al, 2022 [24].

#### Gene Ontology analysis

Biological and molecular GO analysis was performed using *FlyEnrchr*

(https://maayanlab.cloud/FlyEnrichr/).

*Codon Optimality and Translation Efficiency Metrics:*

##### Codon Stabilization Coefficient, CSC

CSC = Pearson’s R (codon frequency_c_, mRNA half-lives)

##### tRNA adaptation Index (tAI)

tRNA adaptation index of a codon, tAI, was calculated using a custom python script according to dos Reis et al., 2004 [27]. Typically, tAI is computed as the weighted sum of the copy number of all *j* cognate decoding tRNAs that read with an anticodon-codon binding affinity s_cj_. Here, we used the mean tRNA expression levels (TMM) to calculate the tissue-specific tAI weights for each codon.

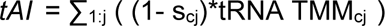

tAI values were amino acid normalized by dividing each tAI_c_ weight by the maximum weight among all synonymous codons of the same amino acid family.

The tAI of a gene, tAI_gene, is the geometric mean of the tAI of all codons in the open reading frame.

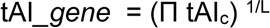

##### Principle Component Analysis

We normalized codon frequencies within amino acid groups to mitigate confounding effects due to amino acid usage and gene length then performed principal component analysis (PCA) followed by unsupervised clustering using Kmeans, where the optimal number of five clusters was determined using the elbow-method.

##### Normalized Codon Frequencies

The length normalized frequencies for all possible 61 codons in a coding sequence (CDS) were computed as follows:

Freq. of codon_i= (number of codon_i)/(total number of sense codons in the CDS) using a custom python package (https://github.com/rhondene/Codon-Usage-in-Python/tree/master/CodonCount).

##### Supply Demand Ratio (SDR)

Codon-level supply-demand ratio (SDR_c_) was calculated as:

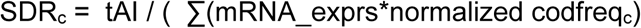

The gene-level supply-demand ratio, SDR, was calculated as the geometric mean of all SDR_c_ values in the coding sequence:

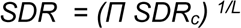

##### Codon Adaptation index

CAI, was computed from coding sequences using DAMBE [39].

#### Hypothesis testing and data visualization

Statistical tests and visualization were performed in R or in python3 (v3.8) using the following packages: scipy, sklearn, pandas, matplotlib, seaborn, statsannotations and statsmodel.

## ACKNOWLEDGEMENTS

We thank Tasos Gogakos and Markus Hafner for hydro-tRNAseq technical guidance and David Ardell for computational advice. We thank Cheng-Yu Lee for sharing *Drosophila* stocks. We also used stocks from the Bloomington *Drosophila* Stock Center (NIH P40OD018537). The authors acknowledge funding from the National Institutes of Health (1R21NS121995 to M.D.C.).

---

**SUPPLEMENTAL TABLE 1.**
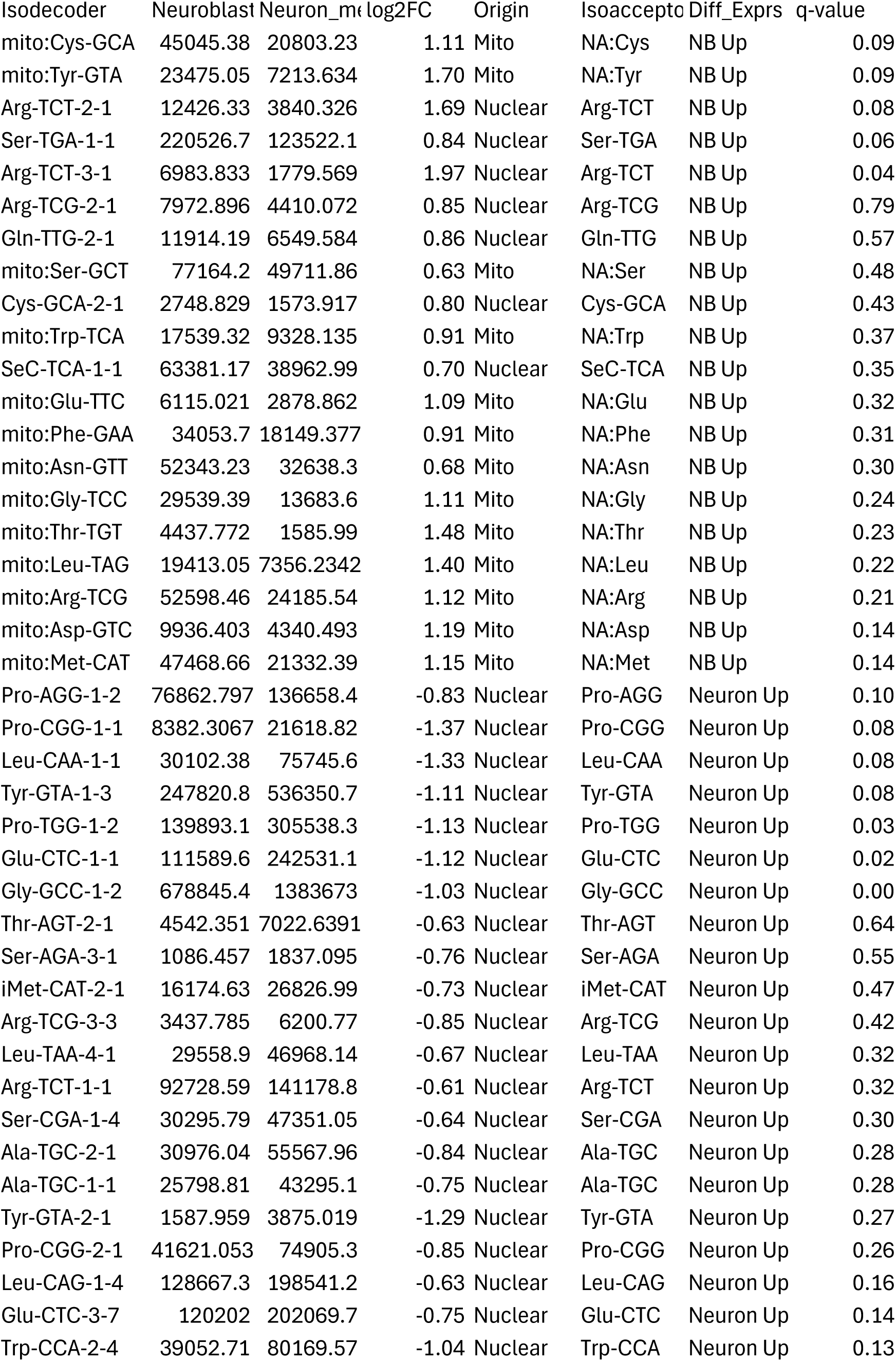
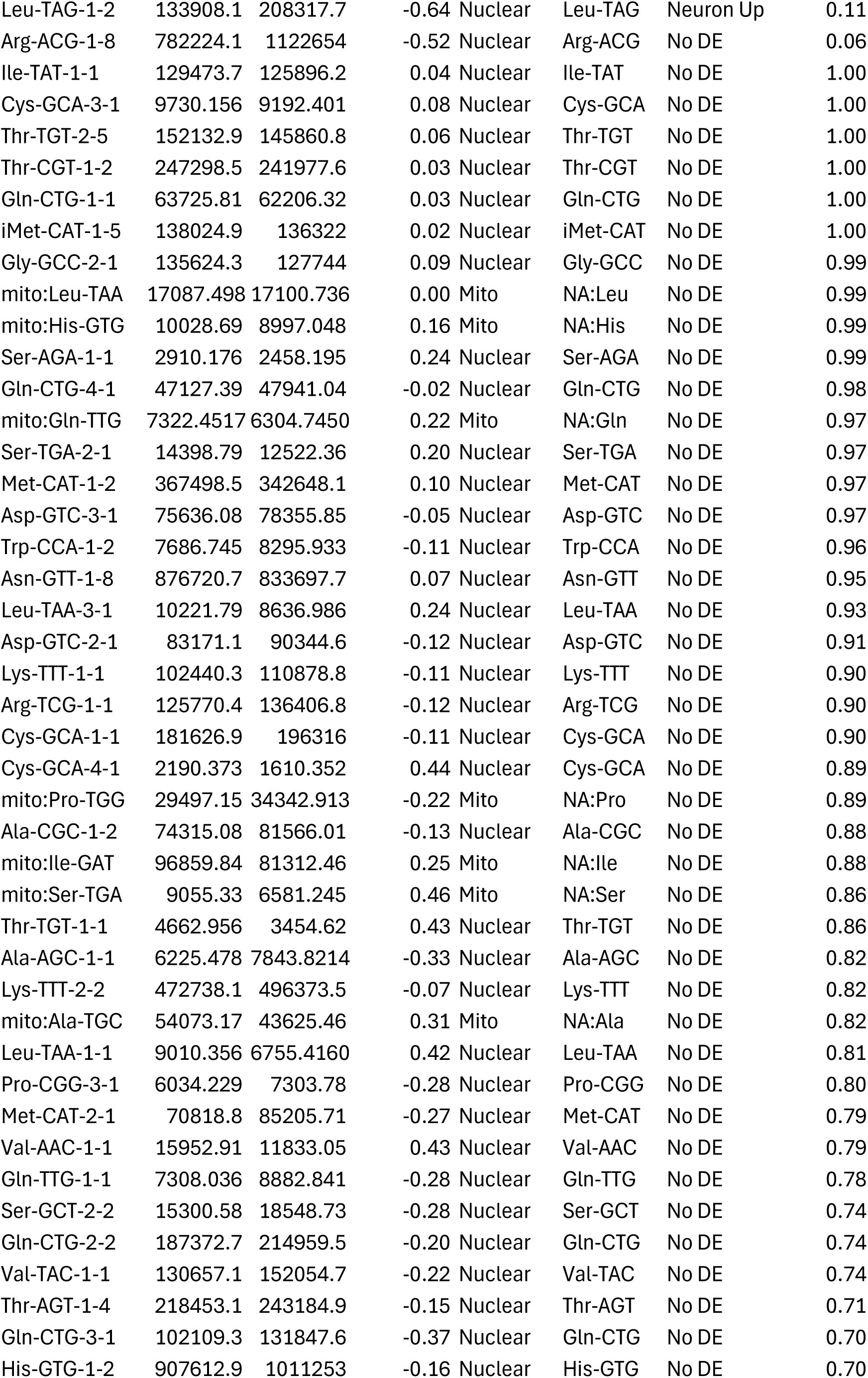
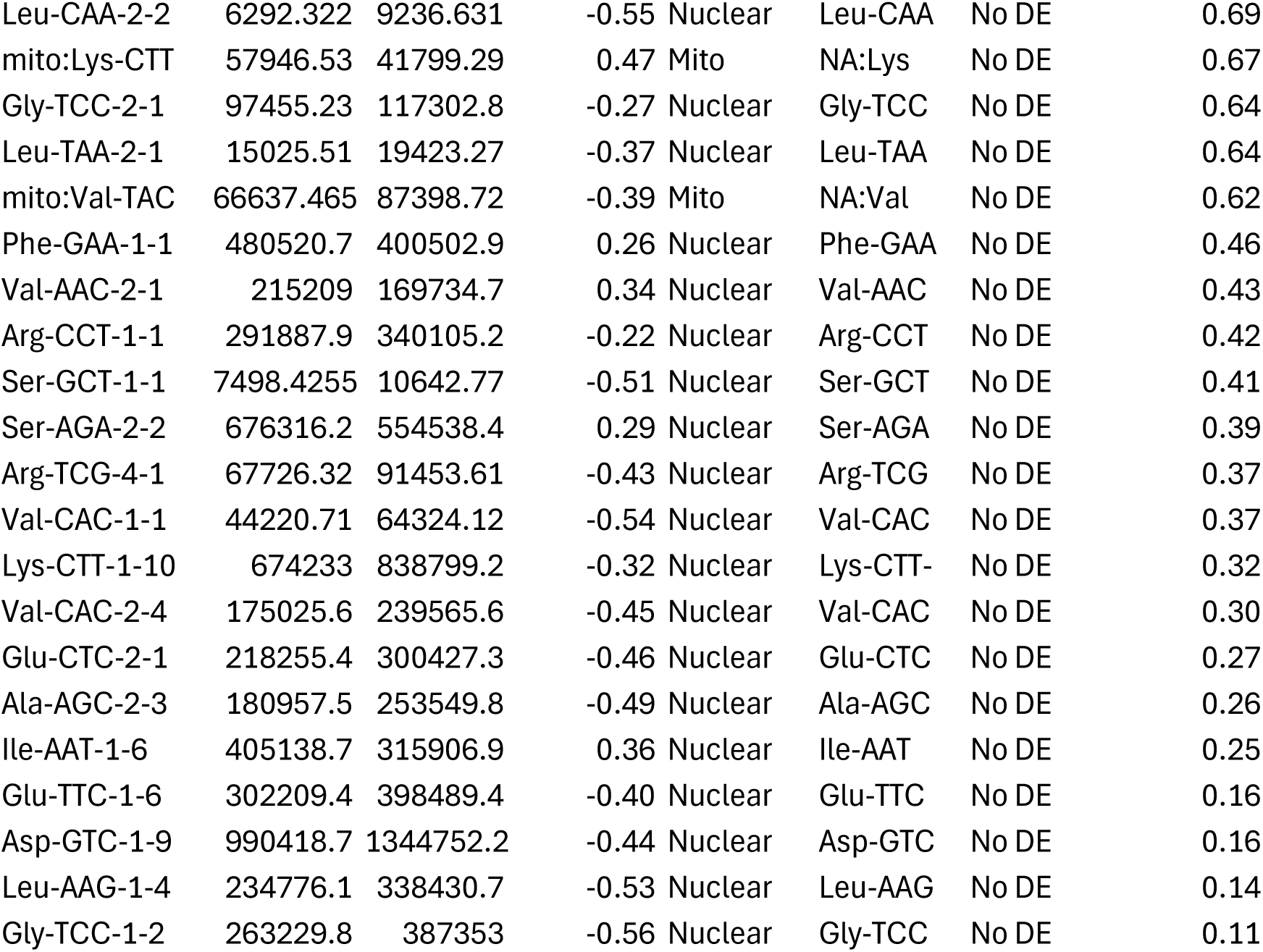

